# scHiMe: Predicting single-cell DNA methylation levels based on single-cell Hi-C data

**DOI:** 10.1101/2022.09.13.507815

**Authors:** Hao Zhu, Tong Liu, Zheng Wang

## Abstract

Recently a biochemistry experiment was developed to simultaneously capture the chromosomal conformations and DNA methylation levels on single cells. A computational tool to predict single-cell methylation levels based on single-cell Hi-C data becomes necessary due to the availability of this experiment. scHiMe was developed to predict the base-pair-specific methylation levels in the promoter regions genome-wide based on the single-cell Hi-C data and DNA nucleotide sequences using the graph transformer algorithm. Promoter-promoter spatial interaction networks were built based on single-cell Hi-C data, and single-cell DNA methylation levels on 1000 base pairs for each promoter were predicted based on the network topology and DNA sequence. Our evaluation results showed a high consistency between the predicted and the true methylation values. We tested using predicted DNA methylation levels on all promoters to classify cells into different cell types, and our results showed that the predicted DNA methylation levels resulted in almost perfect cell-type classification, which indicated that our predictions maintained the cell-to-cell variability. We also tested using the predicted DNA methylation levels of different subsets of promoters and different subsets of CpGs in promoters to classify cells and provided the promoters and CpGs that were most influential in cell-type clustering. Moreover, we observed slightly better performance for the nodes that have higher degree values in the promoter-promoter spatial interaction network but did not find a similar trend for more significant network influencers. Last but not least, we found that using the predicted methylation levels of only housekeeping genes led to less accurate cell-type clustering, which demonstrated that our methylation predictions fit the biological meanings of housekeeping genes since housekeeping genes usually have constant and similar genetic and epigenetic features among different types of cells. scHiMe is freely available at http://dna.cs.miami.edu/scHiMe/.

**Author Summary:** DNA methylation is the process of adding methyl groups to the DNA molecule without changing the nucleotide sequence, which can significantly change the activities of the DNA. Although DNA is a one-dimensional long sequence consisting of cytosine (C), guanine (G), adenine (A), and thymine (T), it folds into a three-dimensional structure in the nucleus of the cell. Scientists believe that this 3D structure has relationships with the activities of the DNA. This research uses deep learning to predict the DNA methylation status of each cytosine-guanine pair in the promoter regions of all human genes based on the 3D structure of the DNA. This not only adds a useful method to the computational biology field but also proves that 3D genome structure does have a relationship with DNA methylation. Moreover, the 3D genome and the DNA methylation are not based on a population of cells, but on individual cells, which provides a cell-specific perspective that allows the understanding of cell-to-cell variability.

## Introduction

DNA methylation in mammals is a biological process in which a methyl group is attached almost exclusively to the cytosine. It regulates gene expression by blocking the binding of transcription factors to DNA or by recruiting the proteins involved in gene repression [1]. Many cellular processes, including embryonic development, X chromosome inactivation, and chromosome stability maintenance, are related to DNA methylation [2]. Reduced representation bisulfite sequencing (RRBS) is the technique that has been widely used to detect DNA methylation levels on bulk cells. Identifying the genomic regions with different methylation levels in bulk cells can differentiate standard and aberrance tissue types [3-5]. In contrast to biological assays, DNA methylation levels of bulk cells can also be predicted by computational models, which save time and effort for wet-lab experimentalists [6-8].

The chromosome conformation capture (3C) technology was developed to reveal the spatial proximities between a single pair of genomic loci in a cell population [9]. After that, 3C-On-Chip (4C) [10], 3C-Carbon Copy (5C) [11], and Hi-C techniques [12] were developed to capture spatial proximities between a locus and all other genomic loci, between all restriction fragments within a given genomic region, and between all possible fragments across the whole genome in a cell population, respectively. Particularly, the Hi-C technique reveals topologically association domains (TADs) as the structural and functional units of the genome [13, 14] and makes it possible to classify TADs into families [15]. The Hi-C data can also be used to computationally reconstruct the 3D structure of the chromosomes [12, 16, 17].

Unlike the standard or bulk Hi-C technique that captures the genome-wide spatial interactions from a population of cells, the recently developed single-cell Hi-C technique can capture the spatial proximities of the whole genome of individual cells and be used to reveal cell-to-cell variability [18]. Like the single-cell Hi-C experiment, the single-cell reduced representation bisulfite sequencing (scRRBS) [19] technique can detect the DNA methylation of single cells. Hui *et al*. [19] used scRRBS to obtain the methylation information of ∼1 million CpG sites of individual diploid mouse and human cells at single-base resolution.

A new experiment protocol named single-nucleus methyl-3C sequencing was recently developed that simultaneously detected both single-cell DNA methylations and the single-cell spatial proximities of the genomes [20]. Moreover, Li *et al*. reported a molecular assay approach named Methyl-HiC that simultaneously captured the chromosomal conformations and DNA methylations in individual mouse cells [21]. However, not all biological wet labs are capable of performing this protocol. Therefore, there are limited data sets available that contain simultaneously-captured single-cell methylation data and single-cell Hi-C data.

The sparsity of the single-cell methylation and Hi-C data is a challenge for computational methods. A frequently-used approach to enhance the data of a single cell is by aggregating the data from other similar cells or building a meta-cell. In the research of [22-24], when a cell has insufficient methylation values, it integrates the methylation data from the neighboring cells. Uzun *et al*. [25] also used the idea of meta-cells when predicting single-cell gene activity levels based on single-cell methylation data. In this research, we applied the meta-cell idea to both single-cell methylation data and single-cell Hi-C data.

In this research, the single-cell Hi-C data are used to build promoter-promoter spatial interaction networks first, and DNA methylations are then predicted based on the network or graph topology.

Learning knowledge or training machine-learning models from graph-structured data has drawn research attention in the computational field because standard neural networks like convolutional neural networks and recurrent neural networks were not designed to work with graph-structured data [26]. In 2017, Monti *et al*. presented the mixture model network [27], a spatial-domain model for deep learning on non-Euclidean domains such as manifolds and graphs, which were applied to different geometric deep learning tasks. Battaglia *et al*. presented the graph networks in 2018 [28], which was a more general framework for learning node-level, edge-level, and graph-level representations. In 2020, the unified message passing model (UniMP) [29] achieved state-of-the-art performance on all the tasks in the open graph benchmark (OGB), a collection of realistic, large-scale, and diverse benchmark datasets for machine learning on graphs [30]. UniMP incorporated feature and label propagation jointly by a graph transformer operator and employed a masked label prediction to optimize the graph transformer. In this study, we modified the architecture of the graph transformer in UniMP to achieve improved performance on our data.

## Results

### Overview

Fig 1 shows the overview of the methodologies used in this research. The single-cell Hi-C data of the target cell was used to generate a promoter-promoter Hi-C contact matrix for the target cell. The idea of meta-cell was then applied, with which the promoter-promoter Hi-C contact matrices of the neighboring cells were aggregated to generate the aggregated promoter-promoter Hi-C contact matrix for the target cell. Based on the aggregated promoter-promoter Hi-C contact matrix, a promoter-promoter spatial interaction network was constructed. The true base-pair-specific DNA methylation values or target values for the 1000 base pairs in the target promoter were generated also based on meta-cell. Node features and edge features were generated and input into the graph transformer network, which contained five blocks of graph transformer. Details of each step will be presented in later sections.

**Fig 1.**
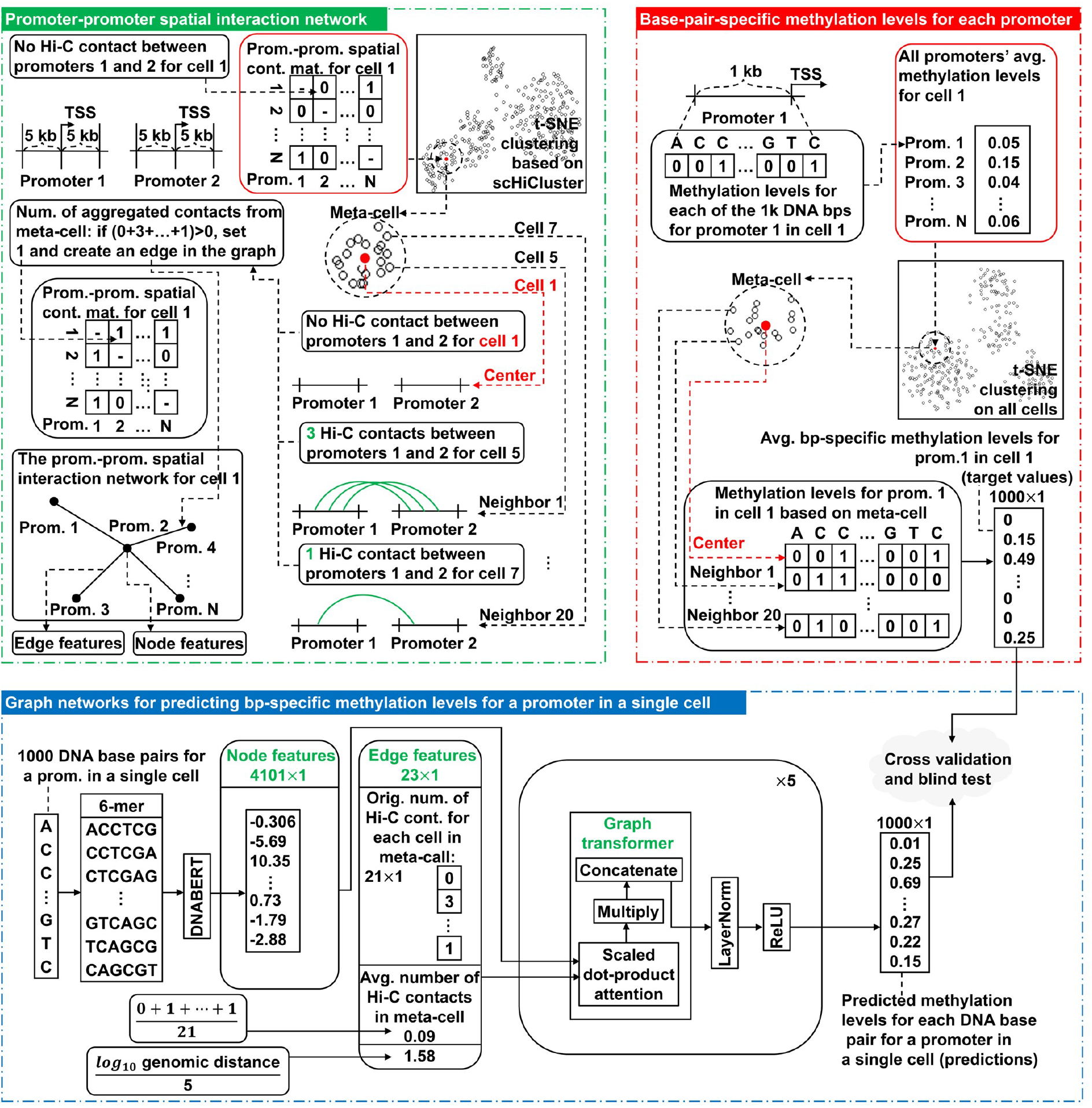
The overview of the architectures of scHiMe. The top left area illustrates the process of building promoter-promoter spatial interaction networks. The top right area shows how the true DNA methylation values, or the target values for the machine learning algorithm, are defined based on meta-cells. The bottom area shows the architecture of the graph transformer including the illustrations of node and edge features.

### Blind test results on data set 1 and data set 2

Table 1 shows the evaluation results for the predictions from scHiMe based on five evaluation metrics, which were conducted on 328 and 1053 cells in the blind-test data sets generated from data sets 1 and 2, respectively. We can find that the overall absolute difference between the scHiMe-predicted and true methylation levels is ∼0.98 on average, and the AUC of the ROC for scHiMe are above 0.92 on both data sets. The performances of scHiMe are much better than the performances of the two naive predictors, which indicates the contributions of the graph transformer in terms of capturing graph topology and data patterns in the networks and improving performances.

**Table1.** The evaluation results for scHiMe and two naive predictors on the blind-test cells on data sets 1 and 2.

| Prediction method | Data set | PCC | Average precision | MCC | AUC | Mean abs. error |
| --- | --- | --- | --- | --- | --- | --- |
| scHiMe | Data set 1 | 0.723 | 0.768 | 0.701 | 0.934 | 0.097 |
| scHiMe | Data set 2 | 0.716 | 0.767 | 0.694 | 0.927 | 0.099 |
| Naive 1 | Data set 1 | 0.059 | 0.189 | 0.058 | 0.533 | 0.286 |
| Naive 1 | Data set 2 | 0.097 | 0.203 | 0.095 | 0.550 | 0.274 |
| Naive 2 | Data set 1 | 0.485 | 0.377 | 0.479 | 0.813 | 0.154 |
| Naive 2 | Data set 2 | 0.545 | 0.441 | 0.530 | 0.836 | 0.148 |

To further analyze the performances of scHiMe, we plotted the absolute differences between the scHiMe-predicted and true methylation levels for different ranges of the true methylation levels, see Figs 2A and 2B for data sets 1 and 2, respectively. We can see that when the true methylation levels are 0 and 1, the absolute differences reach the minimum, which are 0.057 and 0.208 for data set 1, and 0.060 and 0.234 for data set 2, respectively. The largest absolute differences were found when the true methylation levels are in the range of (0.5, 0.6], which are 0.412 and 0.413 for data sets 1 and 2, respectively. However, Figs 2C and 2D show that almost all of the true methylation levels are at 0 and 1, which are 77.9% and 18.5% for data set 1, and 78.9% and 19.3% for data set 2, respectively. The ROC of the scHiMe predictions are shown in Figs 2E and 2F for data sets 1 and 2, respectively.

**Fig 2.**
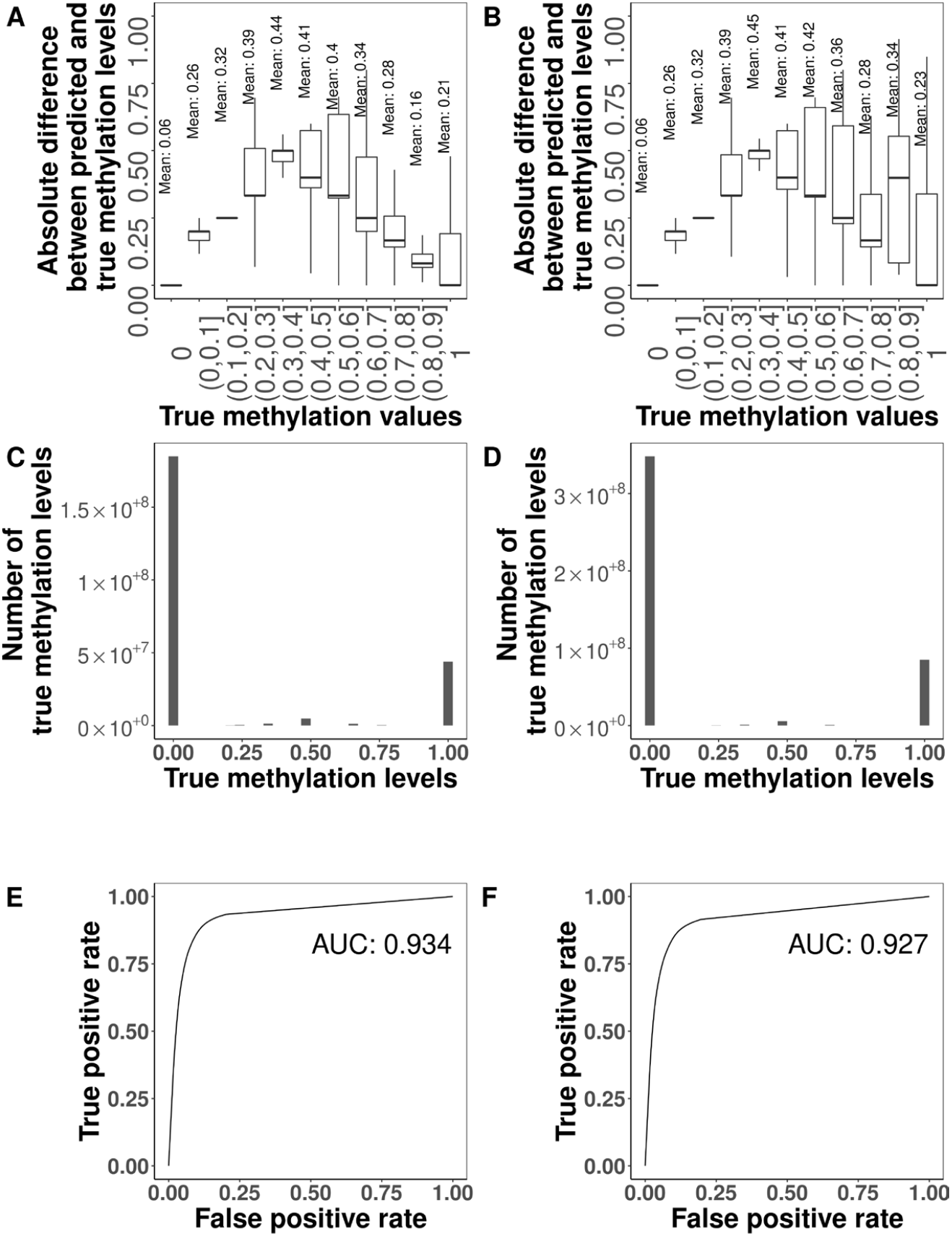
Blind test results on data set 1 and data set 2. (A-B). Absolute differences between the scHiMe-predicted and true methylation levels on the blind-test cells in data sets 1 and 2, respectively. The numbers shown in A and B are the mean values for different ranges of true methylation levels. (C-D) Distributions of the true methylation levels on data sets 1 and 2, respectively. (E-F) ROC curves and the AUC scores for the predictions of scHiMe on the blind-test cells in data sets 1 and 2, respectively.

### Cell-type classification based on the methylation levels of all promoters

Figs 3A and 3B illustrate the first two components from t-SNE based on the true methylation levels on data sets 1 and 2, respectively. For the cytosines and guanines of all CpGs of all promoters, if there are true methylation levels available, the true methylation levels were used for the cytosines and guanines; if not available, −1 was used. It can be found that the cells were clustered almost perfectly to their real cell types, which matched the results of a similar experiment [20].

**Fig 3.**
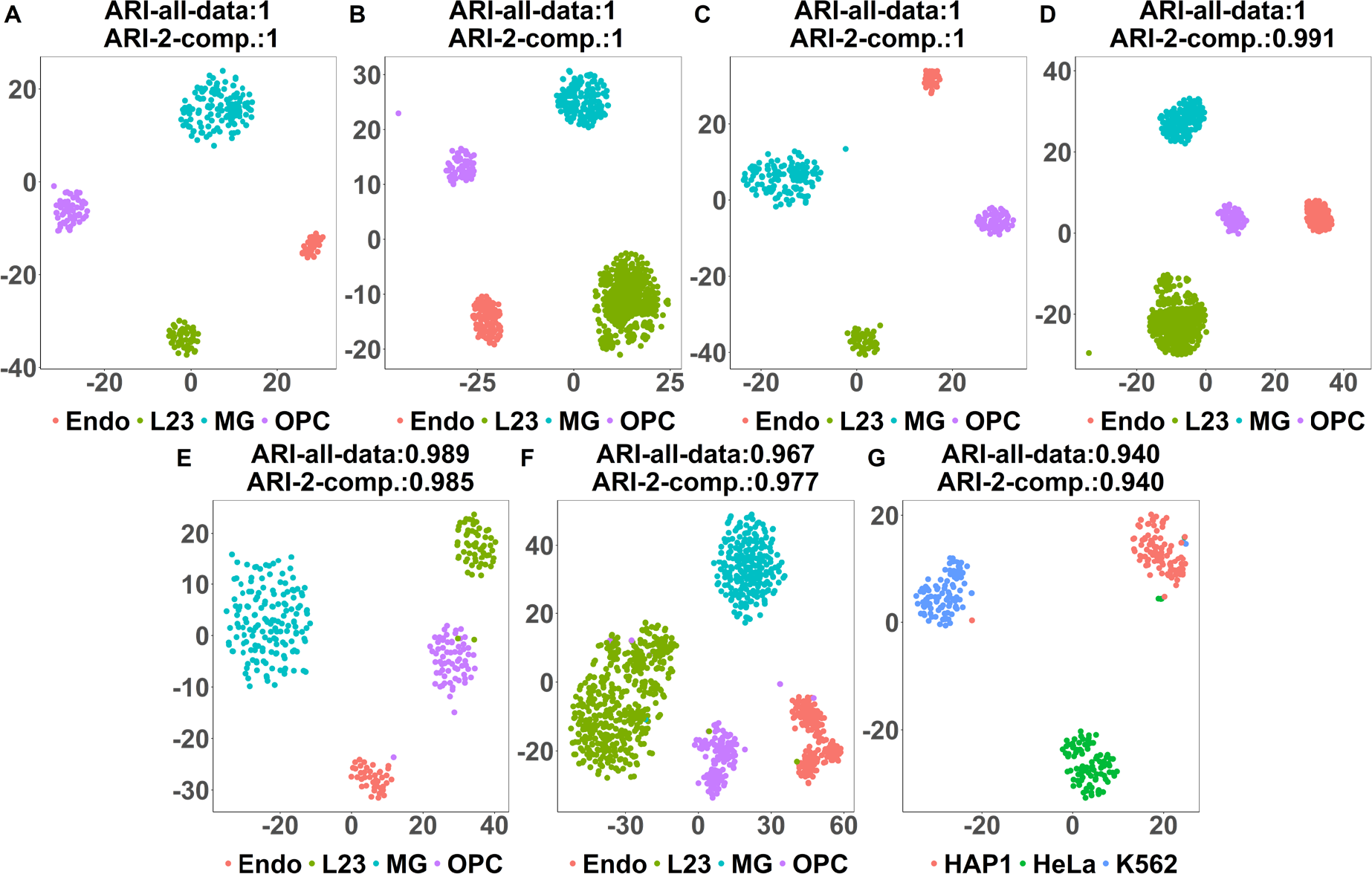
Cell-type classification based on the methylation levels of all promoters. (A) Visualization of the top two components from t-SNE based on true methylation levels on data set 1. For the Cs and Gs of CpGs that did not have methylation data available, −1 was used. (B) Similar to (A), but on data set 2. (C) Visualization of the top two components from t-SNE based on scHiMe-predicted methylation levels on data set 1. For the Cs and Gs of CpGs that did not have methylation data available, −1 was used. (D) Similar to (C), but on data set 2. (E) Visualization of the top two components from t-SNE based on scHiMe-predicted methylation levels on all Cs and Gs of CpGs no matter whether true methylation levels are available or not. (F) Similar to (E), but on data set 2. (G) Similar to (E), but on data set 3.

Figs 3C and 3D are similar to Figs 3A and 3B, but the scHiMe-predicted methylation levels were used for the cytosines and guanines of CpGs that have true methylation levels available. For the cytosines and guanines of CpGs that did not have true methylation levels available, −1 was still used. It can be found that almost all of the cells were still correctly classified. This demonstrates that the predictions from scHiMe maintain the unique patterns of single-cell methylation data for each cell type.

For Figs 3E and 3F, we used the scHiMe-predicted methylation levels for all of the cytosines and guanines of CpGs, no matter whether the true methylation levels were available or not. This experiment tested whether scHiMe-predicted methylation levels resulted in the correct classification of cells even for the cytosines and guanines of CpGs that did not have true methylation data available from the wet-lab experiments. Figs 3E and 3F demonstrate that almost all of the cells were still correctly classified into partitions, which matched their true cell types.

Figs 3G was similar to Fig 3E and 3F, but was on data set 3, which was another independent data set. It can be found that scHiMe achieved similar performance on this data set.

### Cell-type classification based on the methylation levels of individual promoters and individual cytosines and guanines in CpGs

Fig 4A shows the ARI-all-data scores of clustering cells based on the predicted methylation levels of all cytosines and guanines of the individual or selected promoters on data set 2. We used the scHiMe-predicted methylation levels for each promoter to cluster cells and then ranked the promoters based on the ARI-all-data scores. It can be found that using only the first-ranked promoter resulted in an ARI score of 0.664 and using the top 32 promoters brought the ARI score to 0.898. However, using more than top 50 promoters did not further improve the ARI scores. The ARI scores and the visualizations of clustering results on the top two components for the No. 1-200 and top 1-200 promoters can be found in supplementary data shared at http://dna.cs.miami.edu/scHiMe/.

**Fig 4.**
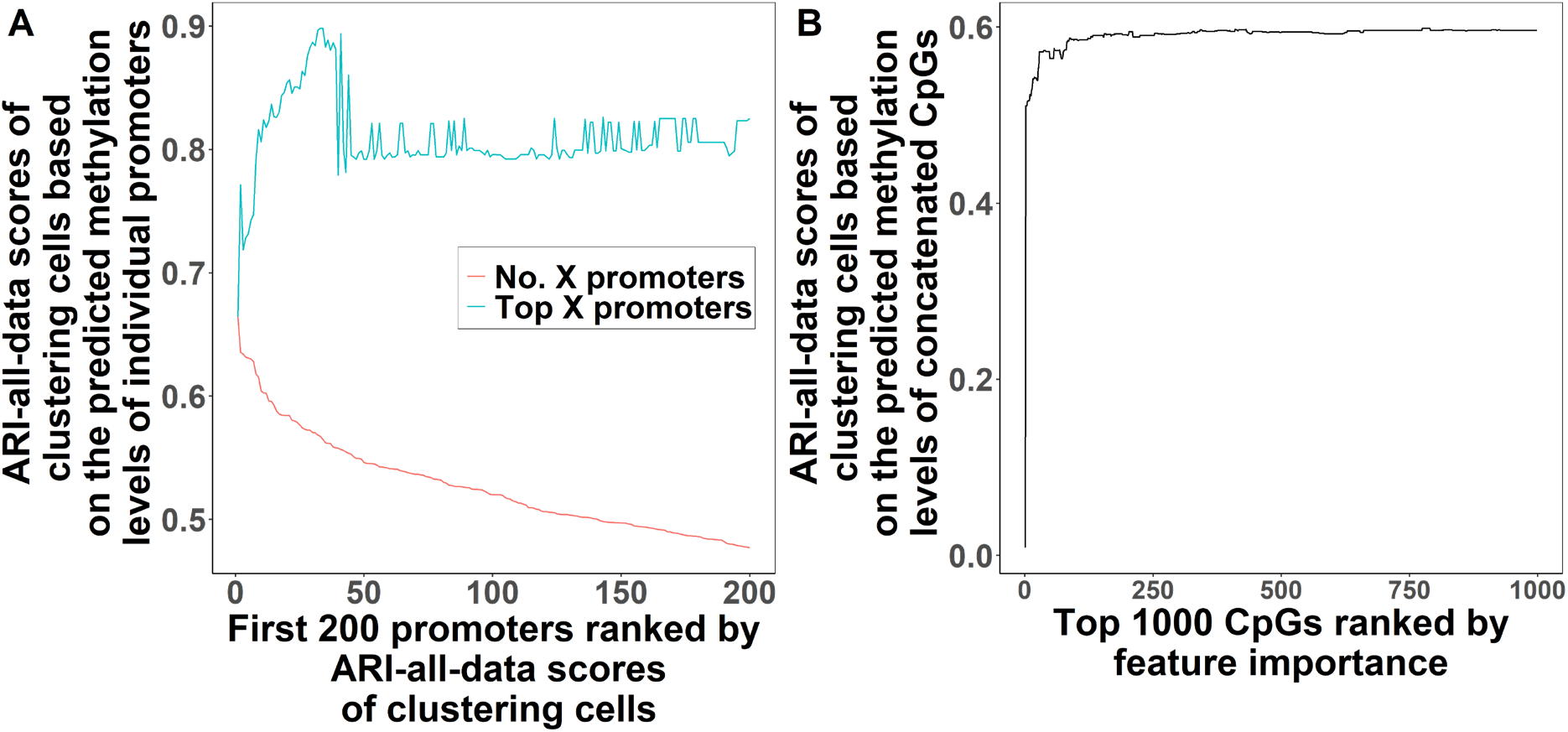
Cell-type classification based on the methylation levels of individual promoters and individual cytosines and guanines in CpGs. (A) ARI-all-data scores of clustering on the first 200 promoters ranked by ARI scores. The first-ranked promoter was labeled as No. 1, the second-ranked No. 2, and so on. The red line indicated the ARI-all-data scores when only No. *n* (1 ≤ *n* ≤ 200) promoter was used, and the values of n were shown as the X-axis. The blue line indicated the performance when top n promoters were used, which meant when all of the first-ranked, second-ranked, …, and the *n*-th-ranked promoters were used. (B) The ARI-all-data scores of the top 1000 CpGs were ranked by feature importance.

We further tested the performance of clustering cells using the scHiMe-predicted methylation level of each cytosine or guanine of a CpG on data set 2. S1 Fig. shows the distribution of the number of cytosines and guanines having various values of feature importance. We ranked all of the cytosines and guanines based on their feature importance and clustered cells using the top n cytosines or guanines. The ARI-all-data scores for the top 1-1000 were plotted in Fig 4B. It can be found that when the top 28 cytosines or guanines were used, the ARI score reached 0.572 and remained there even when the number of cytosines or guanines went up to 1000. The ARI-all-data scores and the visualizations of the top two t-SNE components of the top 1-1000 cytosines or guanines can be found in the supplementary data shared at http://dna.cs.miami.edu/scHiMe/.

### Performance from the perspective of network degree

The degree of a promoter in a promoter-promoter spatial interaction network was defined as the number of edges that connected to the promoter. Fig 5A shows the evaluation metrics for different degree values from 1 to 40 on data set 2. It can be found that the performances are stable when the degree is from 1 to 20, and the performances slightly increase and then vary when the degree is larger than 20. The best AUC 0.965 is with degree 34, and the best Pearson correlation, Matthews correlation coefficient, average precision, and average absolute difference, which are 0.864, 0.853, 0.886, and 0.045, respectively, are with degree 36.

**Fig 5.**
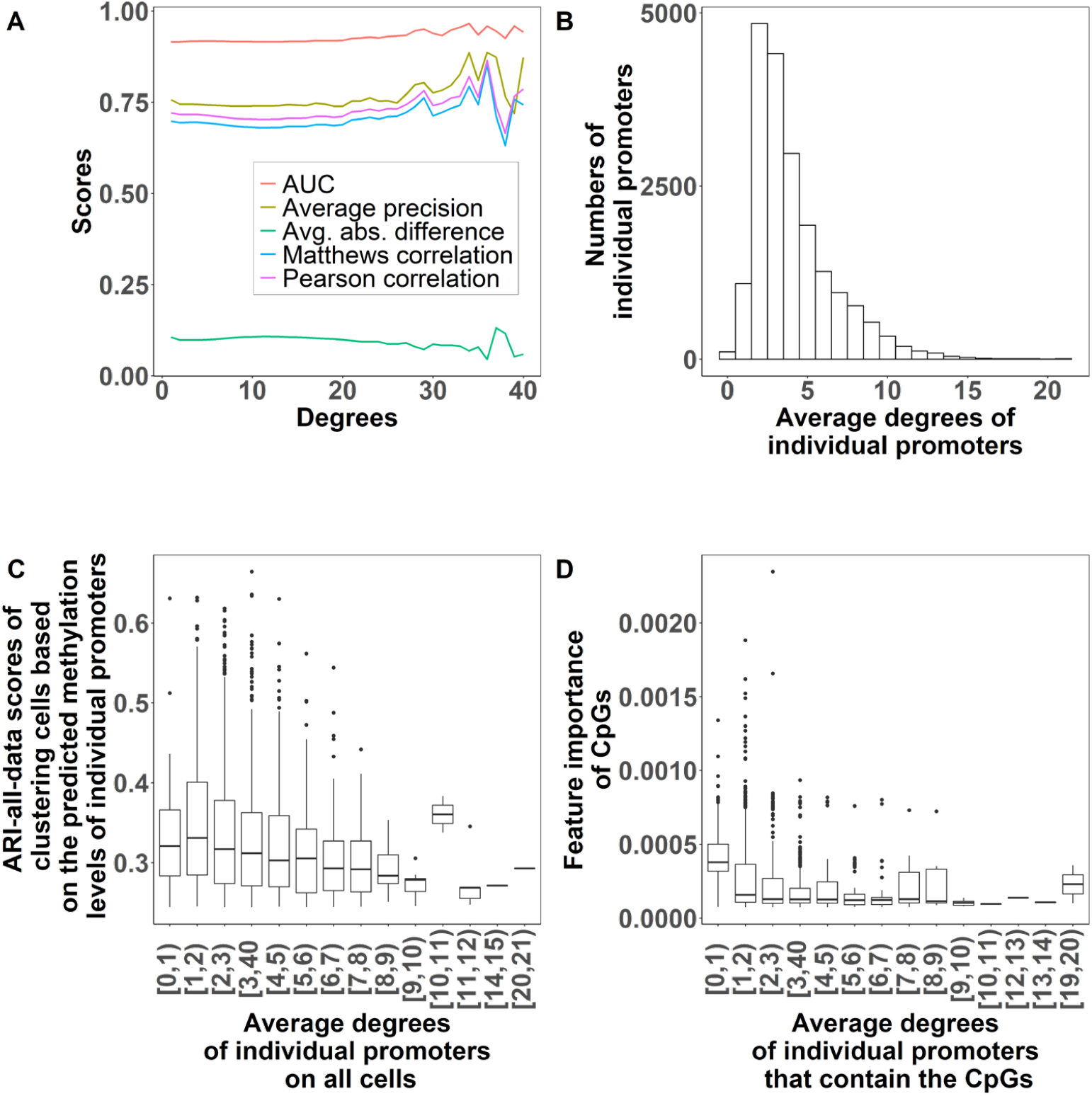
Performance from the perspective of network degree. (A) Evaluation matrices for different degree values on data set 2. (B) Distribution of average degree for all of the promoters on data set 2. (C) ARI-all-data scores of clustering cells based on the predicted methylation levels of the individual promoters of different average degrees. (D). Feature importance for all cytosines and guanines of CpGs from the individual promoters of different average degrees.

Fig 5B illustrates the average degree distribution for all of the promoters on data set 2. We gathered the degree values of each promoter in the promoter-promoter spatial interaction networks of different cells and then calculated an average degree value for each promoter. Fig 5B shows that most of the promoters have average degrees of 2 and 3.

We further tested the performances when using the individual promoters having various average degrees to cluster cells. Fig 5C shows the ARI-all-data scores when using the individual promoters of different average degrees to cluster cells. The promoters used in Fig 5C were the top 3000 promoters ranked by the ARI-all-data scores when each of them was used to cluster cells. It can be found that the promoters with average degrees between 10 and 11 are associated with the highest ARI scores although it needs to be noticed that few promoters fall into the degree range of [10, 11). Other than that, the promoters in the degree range of [1, 2) resulted in slightly higher ARI scores when used to cluster cells into different types. In general, the increase in the average degree values is associated with the decrease in ARI scores. S2 Fig shows the same plot but generated on all promoters, which shows that the promoters with smaller degrees are associated with higher ARI scores.

We gathered the feature importance generated by random forest for all cytosines and guanines of CpGs, and then visualized the feature importance concerning the degrees of the promoters that contained the cytosines and guanines, see Fig 5D. Only the top 3000 cytosines or guanines that have the highest feature importance values were used for the analysis in Fig 5D. Interestingly, the higher the average degree value is, the lower mean feature importance the cytosines and guanines have.

### Performance from the perspective of network influencer

We also benchmarked the performance of scHiMe from the perspective of network influencers. S3 Fig shows the distribution of the number of influencers found for all the cells in data set 2. It can be found that 85% of the cells have from 14007 to 17822 influencers detected from their promoter-promoter spatial interaction networks. We studied the degrees of the top 10 influencers for all cells in data set 2, see the distribution in Fig 6A. We did find higher mean degree values for the first-ranked and second-ranked influencers, which are 7.374 and 6.964, respectively. For the influencers ranked from the third to the tenth, they have similar mean degree values.

**Fig 6.**
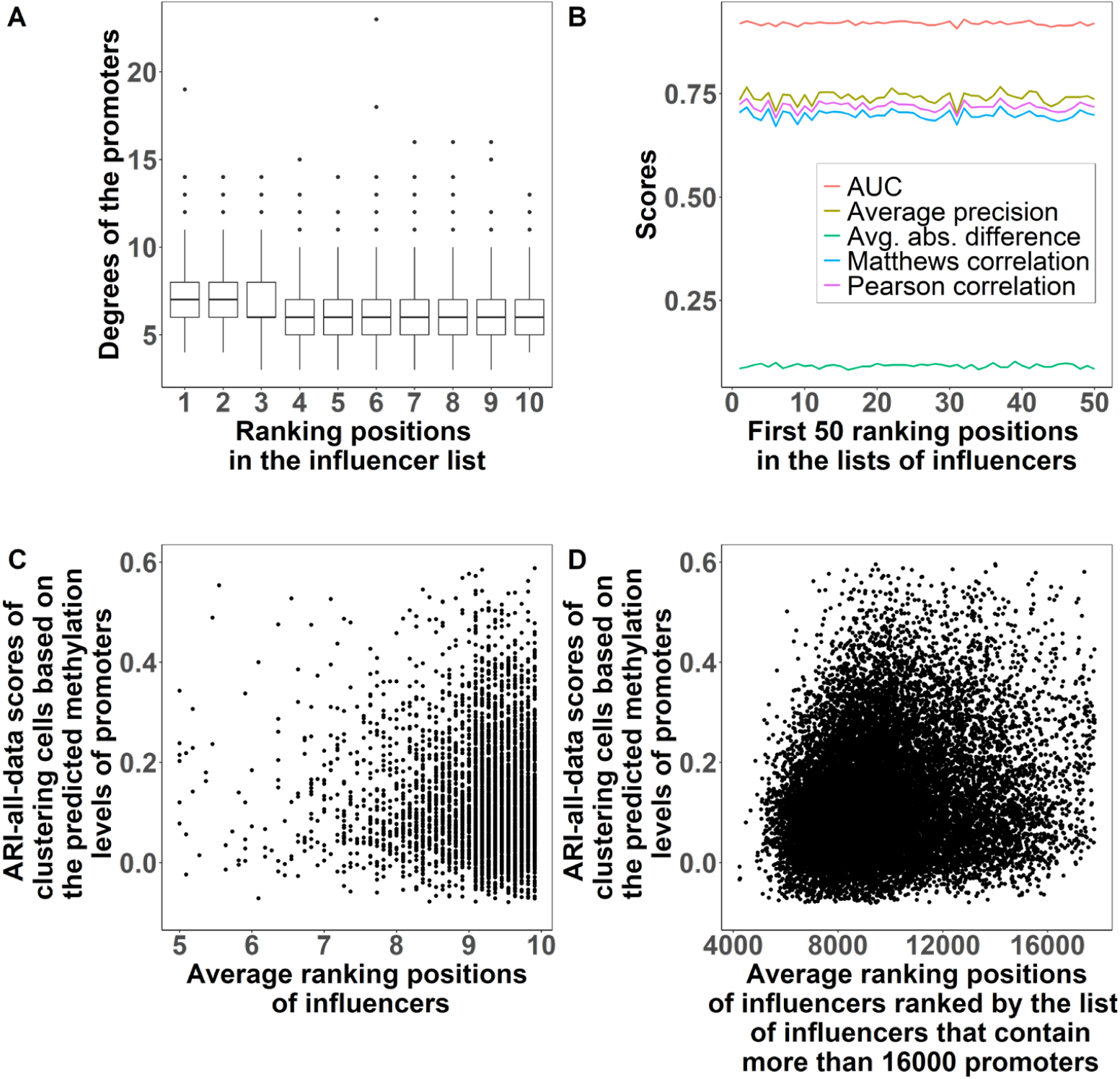
Performance from the perspective of network influencer. (A) The distribution of the degrees of the top 10 influencers for all cells in data set 2. (B) Evaluation matrices for top 50 influencers on data set 2. (C) ARI-all-data scores of clustering cells based on the predicted methylation levels on all cells based on the top 10 most significant influencers (promoters). (D) Similar to (C), but on the cells that have at least 16000 influencers (promoters) in the list.

Fig 6B shows the evaluation metrics for the influencers ranked from the first to the fourteenth on data set 2. We did not find a trend of increasing or decreasing for all of the evaluation metrics with the increase of the influencer ranking positions.

We further investigated the relationship between influencer significance and the feature importance of the cytosines and guanines in CpGs on data set 2, see S4-S5 Figs. Similar to the five evaluation metrics, we did not find an obvious relationship between influencers and feature importance.

We studied whether the most significant influencers resulted in higher ARI-all-data scores when the predicted methylation levels on the influencers were used to cluster cells into different cell types. Fig 6C shows our benchmark results on all of the cells but only considers the top 10 most significant influencers. Fig 6D illustrates the results on the cells that have at least 16000 influencers detected, which has a Pearson correlation of 0.098 between influencer ranking positions and ARI-all-data scores. In summary, the results from Fig 6C and 6D do not show a strong correlation or relationship between the influencer ranking positions and ARI scores. The average ranking positions in Fig 6C and 6D were calculated using avg_inf_rank1. S6 Fig shows the distributions of the number of influencers with various avg_inf_rank1 values associated with Fig 6C and 6D, respectively. S7 Fig shows the results using avg_inf_rank2 on data set 2, which indicates the same conclusions.

Last but not least, we picked up the influencers that were first-ranked in the influencer lists of all cells and counted the number of occurrences of each promoter as the first-ranked influencer among all cells. The promoters that have the same number of occurrences were used to cluster cells, and the ARI-all-data scores were calculated. Besides the first-ranked influencers, we also conducted the same experiments for the second-ranked, third-ranked, fourth-ranked, and fifth-ranked influencers, see S8A-S8E Fig. No significant Pearson correlation or relationship was found between the ARI-all-data scores and the number of occurrences. We also picked up the top five most occurred influencers for influencer ranking positions 1 to 93 (the shortest influencer list contained 93 influencers) and used the top five influencers to clustering cells, see S8F Fig, from which the same conclusion was drawn.

### Cell clustering and feature importance for housekeeping genes

Housekeeping genes are the genes that are required for maintaining basic cellular functions, which are essential for the existence of cells. Housekeeping genes usually are considered to be expressed in all cells with stable expression levels. Therefore, if the scHiMe-predictions are accurate, the predicted methylation levels on the promoters of housekeeping genes should be less effective in distinguishing cell types compared to the promoters of baseline genes. Fig 7A shows the ARI-all-data scores when housekeeping genes and randomly-selected genes were used to classify cells. The mean of ARI scores for using housekeeping genes is 0.128, which is lower than the ARI score for using randomly-selected genes, which is 0.146, with a P-value of 0.0028. Fig 7B shows the feature importance of cytosines and guanines of CpGs when housekeeping genes and randomly-selected genes are used, which have the means of 1.542×10^−5^ and 1.601×10^−5^, respectively, with a P-value of 0.0059. These results further proved that the predictions from scHiMe matched biological meanings.

**Fig 7.**
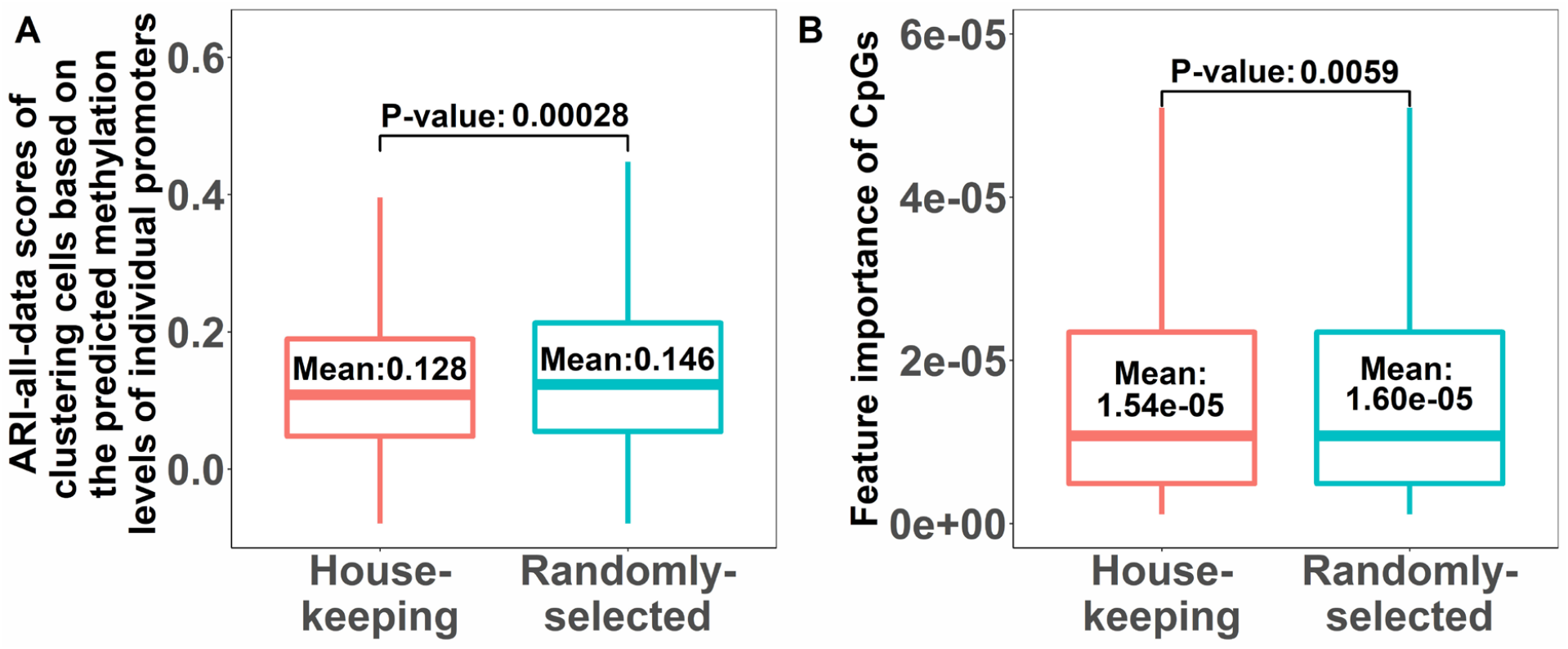
Cell clustering and feature importance for housekeeping genes. (A) ARI scores of using scHiMe-predicted housekeeping genes and randomly-selected genes to classify cells. The P-value between the ARI scores in the two groups is shown in the figure. (B) Feature importance of the cytosines and guanines of the promoter CpGs for housekeeping genes and randomly-selected genes when their scHiMe-predictions were used to classify cells. The P-value between the ARI scores in the two groups is shown in the figure.

## Discussion

We developed a computational tool named scHiMe to predict the single-cell methylation levels of cytosines and guanines of the CpGs in promoters based on single-cell Hi-C data. Graph transformer, one of the cutting-edge deep learning algorithms, was applied. We benchmarked the predictions of scHiMe from multiple perspectives and found that the predictions were accurate and maintained cell-to-cell variability. The DNA sequence used for all of the cells was the same since we used the reference genome to generate features. Therefore, if we only used DNA sequences as machine learning features, the predictions on all the cells would be the same, which meant the cells would not be correctly classified into cell types. However, our predictions resulted in almost perfect cell-type classifications, which indicated that the single-cell Hi-C data that we used indeed provided useful information for cell-specific characteristics, the graph transformer that we applied indeed successfully captured the patterns in the graph-structured data, and our predicted DNA methylations keep the cell-to-cell variability. Moreover, the successful performance of our tool indicated that the single-cell 3D genome structure does have an influential relationship with single-cell DNA methylation, although, from this single study, we were not certain about which influences which or how one of them impacts the other.

## Materials and Methods

### Data sets

The first dataset used in this research contains the simultaneously-captured single-cell Hi-C data and DNA methylation data from the brain cells of a 29-year-old male caucasian [20]. The cells in this data set are from 14 different cell types., and we only kept and used nine cell types (Astro, ODC, Ndnf, Sst, Vip, Endo, L23, MG, and OPC) that contained at least 30 cells. We used the cells in the Astro, ODC, and Ndnf cell types to build the training data set, cells in the Sst and Vip cell types for the validation data set, and the cells in Endo, L23, MG, and OPC cell types for the blind-test data set. We also applied the same quality control as in research [20], based on which we only kept the cells that had total non-clonal reads > 500,000, global mCCC < 3, total autosomal cytosines covered < 100M, and total long-range (>10,000 bp) cis contacts > 5000. Additionally, the total long-range cis contacts of each chromosome need to be larger than x, where x is the length of a chromosome at Mb resolution. This first dataset was used for training, validation, and testing and was also used to benchmark the performance of classifying cells into different cell types based on the predicted single-cell methylation levels of promoters.

The second dataset is similar to the first but from a 21-year-old male caucasian also from the research [20]. We picked the same four cell types: Endo, L23, MG, and OPC and applied the previously-mentioned quality-control criteria. This dataset was only used for blind testing and clustering cells into cell types based on the predicted single-cell methylation levels of promoters.

We used another human single-cell Hi-C data as our third dataset that was from [31]. We used three cell types: K562, HAP1, and HeLa, and selected 100 cells from each cell type that had the highest number of single-cell Hi-C contacts. Since single-cell methylation data were not included in this data set (the authors only performed single-cell Hi-C experiments), we only used it for benchmarking the performance of classifying cells into cell types based on the predicted methylation levels of promoters.

The transcription start site (TSS) definition used in this research was downloaded from the GENCODE human annotation version 19 [32].

### Building meta-cells for single-cell Hi-C data

Due to the sparsity of the single-cell Hi-C data of a target cell, we combined the target cell and other cells, which share similar Hi-C patterns with the target cell, into a meta-cell and used the single-cell Hi-C contacts of the meta-cell as the Hi-C contacts for the target cell. For the cells used for training and validation, their cell types were considered known information. Therefore, the similar or neighboring cells used to build each meta-cell were selected from the same cell type as the target or central cell.

For each chromosome of every cell, we built a promoter-promoter contact matrix based on the single-cell Hi-C contacts of that cell (not meta-cell). Each promoter region here was defined as containing 50 kb upstream and 50 kb downstream of the TSS plus the TSS of a gene. Notice that a promoter was defined as a 1 kb region upstream of a TSS for methylation meta-cell and prediction (details will be discussed in later sections), and here we made each promoter region larger. This is because, in this way, more single-cell Hi-C contacts can fall in the promoter regions. In other words, having the regions too small will result in few Hi-C contacts between the regions, which will not adequately define the graph topology of the promoter-promoter interaction networks. A contact was considered to exist between two promoters if one end of a Hi-C read-pair was located inside a promoter region and the other end located inside another promoter region.

We performed the incremental principal component analysis function (IPCA) using the sklearn package [33] on the flattened promoter-promoter contact matrix of each chromosome and then obtained the top 30 principal components. We tested different number of principle components and found 30 resulted in the minimal explained variance, see S9 Fig. After concatenating the 30 principal components of 23 chromosomes (Y-chromosome not included as it contains only 63 promoters and most of the promoters do not have single-cell methylation data available), we obtained 690 principal components for each cell. We repeated the IPCA on the 690 principal components and further obtained another top 30 principal components. After that, we used these 30 components to compute the Euclidean distance between each pair of cells. Finally, we picked up the 20 nearest neighboring cells and then combined them with the central cell to generate the meta-cell for the target cell. The Hi-C contacts in the meta-cell were used to define edges in the promoter-promoter spatial interaction network.

For the cells used for blind test or benchmarking, we considered the type of each cell unknown, but the number of total cell types in the testing data set was known because most of the biological labs would have known how many and what cell types they used in their experiments. Because of these, we clustered all the cells in the test dataset and then generated the meta-cell for each cell only using the other cells in the same cluster. We executed scHiCluster [34] on the intrachromosomal Hi-C contact pairs of all the cells at 1 Mb resolution. The results of scHiCluster are *n* × (*n* − 1) components, where *n* is the number of cells and *n* − 1 is the number of components for each cell.

We used the spectral clustering function from the sklearn package [33] to cluster all the cells by labeling the *n* cells based on the output components of scHiCluster with the arguments of affinity = nearest_neighbors and cluster = 4. We used all of the components that output from scHiCluster to compute the Euclidean distance between all cell pairs. After that, we selected the 20 nearest neighboring cells in the same cluster of the target cell and then combined the Hi-C contacts of these 20 neighboring cells with Hi-C contacts of the target cell to generate the aggregated Hi-C contacts for the target cell.

### Building meta-cells for single-cell DNA methylation data

Due to the sparsity of the single-cell methylation data, we combined the target cell with other neighboring cells, which share similar methylation patterns with the target cell, into a meta-cell and used the aggregated single-cell methylation levels of the meta-cell as the methylation levels for the target cell.

The single-cell methylation data that we have used provide the number of methylated reads and the total number of reads for individual CpG. We calculated an overall methylated ratio for each promoter of a cell by dividing the sum of the methylated reads (only on CpGs) available for the promoter by the total number of reads (only on CpGs) available for the promoter. Some of the promoters in some of the cells have no methylation data at all. That is, none of the CpGs in the promoters have any reads available, or the promoters do not contain any CpG. For those promoters, we computed the mean overall-methylated-ratios based on all of the other promoters from the same cell and used that mean value as the overall methylated ratio for each of those promoters that did not have methylation data available.

We calculated a variance value for each promoter based on the overall methylated ratios of the promoter in all cells. After that, we selected 5000 promoters with the highest variance values and performed IPCA on their overall methylated ratios. For each promoter, we obtained the top 30 components, which were used to calculate the Euclidean distance between every cell pair. Based on the Euclidean distances, we chose the 20 nearest neighboring cells and then combined them with the target cell to generate the meta-cell.

The aggregated methylation levels of the meta-cell were used as the true or target methylation levels of the target cell. For example, for base pair 1 in the promoter *p*_1_, five of the 21 cells in the meta-cell have methylation data available. Each of the five cells provides both the number of methylated reads on that base pair and the total number of reads on that base pair. The true methylation level or the target value for base pair 1, which was a value in the range of [0, 1], was calculated as the total number of methylated reads from the five cells divided by the total number of reads from the same five cells. If none of the 21 cells in the meta-cell had methylation levels available on a base pair, then we labeled −1 as the target value for the base pair. If a nucleotide base in a promoter was not the cytosine (C) of a CpG, we used −1 as the target value for the base pair unless the nucleotide base was guanine (G) and its complementary base was the cytosine (C) of a CpG. In that case, we used the methylation level of the complementary base as the target value of the base pair.

In summary, each promoter has 1000 target values, one for each base pair in the promoter region.

Notice that the final predictions from scHiMe also contain 1000 values for each promoter (one for each base pair), and the predictions are provided based on the reference genome, which is the DNA sequence of the positive strand. Therefore, the predicted methylation level on a G of a CpG dinucleotide can be considered as the prediction made for its complementary C in the negative strand.

### Promoter-promoter spatial interaction network

In the promoter-promoter spatial interaction network, each node represents a promoter. The aggregated Hi-C contacts from meta-cells were used to define edges in the promoter-promoter spatial interaction networks. An edge was created between two promoters if at least one aggregated Hi-C contact exists between the two promoter regions. The promoter region used here was defined as the genomic region that contains 5 kb upstream and downstream of the TSS plus the TSS of a gene.

### Node and edge features

The node features were the encodings of DNA nucleotide sequence for each 1-kb promoter/node, which was generated by DNABERT [35]. Specifically, we generated the 6-mer representation for each base pair in the DNA sequence and input the representations of all base pairs into DNABERT, which outputted 4101 real numbers as the codings of the promoter. For the promoters existing in the negative strand, we used their complementary sequences in the positive strand in the order of 5’ to 3’ (this 5’ to 3’ is in terms of the positive strand) as the sequences that were input to DNABERT.

The first edge feature contains 21 integers, which are the numbers of Hi-C contacts, between the two promoters that the edge connects in the target cell and 20 neighboring cells in the meta-cell. The second edge feature is the average number of occurrences of nonzero Hi-C contacts in the meta cell.. For example, if there are only two nonzero Hi-C contact values among 21 Hi-C contact values in the meta-cell, which are 3 and 1, as shown in the example in Fig 1, then the second edge feature is 0.095, which is calculated as 2 / 21. The third feature is calculated as 1 / 5 of the base-10 logarithm of the genomic distance between the two promoters that are connected by the edge. The genomic distance here is at a resolution of 1 bp. In total, the edge feature for each edge consists of 23 values.

### Graph transformer

The node features that are input to the o-th block of the graph transformer are 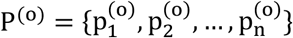, where p stands for a promoter, and n is the number of promoters. Based on the node features, a query vector for the i-th node feature is defined as: 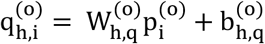, where h indicates that it is the h-th head in the block, and 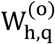 and 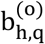 are the trainable parameters for the query vector.

Similarly, a key vector for the j-th node feature is defined as: 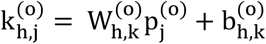.

The edge features input to the transformer block are represented as 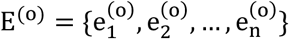.

Specifically, the edge feature for the edge between node i and node j after the h head-attentions is encoded as: 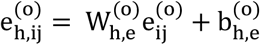, where 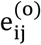 the original edge features between node i and node j, and 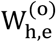 and 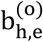 are trainable parameters.

The encoded edge features will be added to the key vector for calculating the multi-head attention: 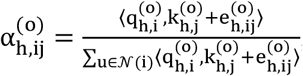, where ⟨q, k + e⟩ is the exponential scale dot-product: 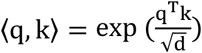, where d is the hidden size of each head attention.

After the above h-th head attention is completed, a value vector is defined as: 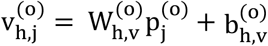, where 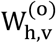 and 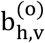 are trainable parameters.

Message aggregation is performed to integrate the value vectors of all the radius-one neighboring nodes, each of which is labeled as j, to node i as: 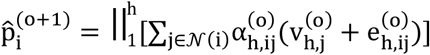, where 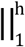 is the concatenation operation that is performed on all h head attentions.

Different from the paper [29], we directly applied the normalization layer and ReLU layer to the multi-head attention as follows, which, based on our evaluation, resulted in better performance (results of comparisons not shown): 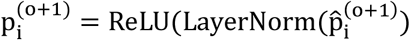.

Also, different from the paper [29], which only aggregated head attention for the node features, we also aggregated the head attentions for the edge features as follows, which resulted in better performance (results of comparisons not shown): 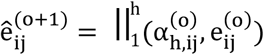.

A normalization and ReLU layer are added to get the edge features that will be input to the next block: 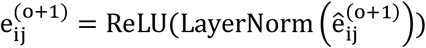.

For the last block, the ReLU layer and the normalization layer are not applied. Instead, the concatenation operation on all h head attentions is replaced by the average on multi-head output as: 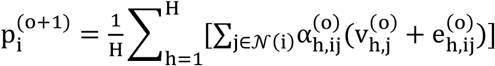.

### Training, validation, and blind test

Three cell types in the first data set were used as the training data, which were Astro, ODC, and Ndnf cell types. The total number of graphs used for training is 21,745. The graph transformer that we used was modified based on the transformer_conv in the torch_geometric.nn.conv pack [36], which was published by [29]. We tested different values of the number of transformer blocks, including 3, 5, 7, and 10, and different values of head attentions including 1, 2, and 5, and found the number of blocks of 5 and the number of head attentions of 1 resulted in the best performance, which were used in our final model.

We used the Adam algorithm [36] as our optimizer to train the graph transformer. We used the function ReduceLROnPlateau in the torch.optim package [36] with the parameters mode=min, patience = 5, and factor = 0.1 to automatically reduce learning rates. The training loss was calculated as the average absolute differences between the predicted methylation levels and the target or ground-truth methylation levels. The training loss was calculated on the base pairs that did not have −1 as the target values.

We used the Vip and Sst cell types from the first data set to generate the validation data. There are 2254 graphs for validating the graph-transformer model. We terminated the training process when the validation loss stopped decreasing for five consecutive epochs or the number of epochs reached 100. The model with the smallest validation loss was selected as the final model and benchmarked on the blind test data sets.

We used Endo, L23, MG, and OPC cell types from the first and second data sets, which corresponded to 7544 and 24219 graphs, respectively, to generate the blind test data. From the third data set, we also chose the top 100 cells having the highest number of Hi-C contacts from each of the K562, HAP1, and HeLa cell types to create the extra blind test data.

### Evaluation

#### Naive predictors

Since there is no other published tool to compare performance, we built two naive predictors. The predictions for a promoter from naive-predictor-1 are the average methylation levels gathered from the radius-one neighboring promoters in the promoter-promoter spatial interaction network. For naive-predictor-2, we defined a target region for each of the base pairs in the target promoter, which contains the target base pair, 50 bp upstream of the target base pair and 50 bp downstream of the target base pair. All of the single-cell Hi-C contacts with one end falling into the target region were collected. For each of the other ends in these Hi-C contacts, we defined an in-contact region, which contains the in-contact base pair that formed the Hi-C contact with the target base pair, 50 bp upstream of the in-contact base pair, and 50 bp downstream of the in-contact base pair. An average methylation value was calculated based on the available ground-truth per-base-pair methylation levels within all of the in-contact regions, which was considered the final prediction from naive-predictor-2 for the target base pair.

#### Performance metrics

To benchmark the performance of scHiMe, we calculated the Pearson’s correlation (PCC), Matthews correlation (MCC), average precision (AP), and area under the curve (AUC) of the receiver operating characteristic curve (ROC) between the predictions and ground truths. Our evaluations were performed only on the base pairs that had true methylation levels available.

We parsed the predicted methylation levels on all the cytosines and guanines in CpGs because guanine in the positive strand indicates a cytosine in the negative strand, and our predictions are on base pairs, not nucleotide bases.

#### Cell classification and its evaluation

The predicted methylation levels on all the CpGs of all promoters were input into the K-means clustering algorithm with the number of clusters set to four. To evaluate the clustering results or evaluate whether the groupings of cells in the clustering results match the true cell types, we applied the adjusted rand index (ARI) to measure the similarities between the clustering results and the true cell types. We named this type of ARI score ARI-all-data. A close to zero ARI score indicates random labeling, and the ARI score of one indicates that two clusterings are identical.

To visualize the predicted methylation levels on all the CpGs of all promoters, we input the data into the t-distributed stochastic neighbor embedding (t-SNE) algorithm, and the top two components were obtained, which were treated as the X and Y coordinates for plotting the cells in a two-dimensional (2D) space. The cells within each true cell type were illustrated by a unique color for visualization. The top two components from t-SNE were input into the K-means clustering algorithm with the number of clusters set to four. Another type of ARI score was calculated based on the clustering results, which were referred to as ARI-two-components in this paper.

Besides clustering cells using all promoters, we also benchmarked the performances of clustering cells using individual promoters or individual CpGs because we believed this provided biologists with more information about which promoters or CpGs and their methylation levels were important in differentiating cells to different cell types. In these cases, the same methodologies for clustering, calculating ARI scores, and visualization were applied with the only difference that the methodologies were applied on the methylation levels of one or selected promoter(s), cytosine(s), or guanine(s).

#### Importance of individual CpGs in promoters for clustering cells

We used the random forest algorithm from the sklearn package [33] to measure the importance of individual CpGs in promoters for classifying cells. Random forest was used to classify a cell into one of the four cell types: Endo, L23, MG, and OPC. We used the predicted methylation levels on all of the CpGs, including both cytosines and guanines, from all of the promoters genome-wide as the features. In other words, each training example is for one cell with the target value as the true cell type for that cell, followed by 1443574 predicted methylation levels. We used all of the 1053 cells in the four cell types in data set 2 to generate the training and validation data set for the random forest: 80% of the cells were used for training and 20% for validation. We tested different values for the number of trees, including 100 and 1000, which resulted in classification accuracies of 0.857 and 0.995, respectively. Based on these, the feature importance was generated based on the random forest model that consisted of 1000 trees. Each of the cytosines and guanines in all of the CpGs had a feature importance value output from the random forest algorithm, which indicated the importance of each cytosine or guanine when used to classify cells into the four cell types.

#### Cell clustering based on housekeeping genes

Housekeeping genes are typically constitutive genes that are required for the maintenance of basic cellular function. We downloaded the human housekeeping genes from the HRT Atlas database [37]. Among all the housekeeping genes that we downloaded, 1967 of them matched the genes that were used to define the promoters in our research and were used in this analysis. We randomly selected another 1967 promoters from all of the promoters that we had in the research and used them as the baseline analysis.

#### Collective Influence algorithm

We applied the collective influence (CI) algorithm [38] to the promoter-promoter spatial interaction networks built on data set 2. A promoter-promoter spatial interaction network was built for each chromosome of a cell, and all of the promoter-promoter interaction networks of a cell were saved in one file and input into the CI algorithm for finding network influencers. The nodes that had no edge connecting to any other nodes were removed. A list of influencers ranked from the most significant to the least significant influencers was outputted from the CI algorithm for each cell.

To find the relationship between the ranking position of each promoter in the influencer list and the ARI score of using the promoter to classify cells, we defined two ways of calculating the average ranking position in the influencer list for each promoter, which are named avg_inf_rank1 and avg_inf_rank2. For each promoter in avg_inf_rank1, if its ranking position for a cell was smaller than *t*, then this original ranking position was used. If its ranking position for a cell was larger than *t*, or if the promoter was not even included in the ranking list for a cell, then *t* + 1 was used as its ranking position for that cell. Two values for threshold *t* were tested, which were 10 and 17822. When using 10, we focused on the cases where a promoter was among the top 10 most significant influencers.

When using 17822, we included all possible ranking positions since the longest ranking list contained 17822 influencers. When using the threshold 17822, we only used the cells that had at least 16000 influencers output from the CI algorithm.

For each promoter in avg_inf_rank2, if its ranking position for a cell is smaller than *t*, then this ranking position was kept. We gathered all of the ranking positions for the promoter that meet this criterion and then calculated the average ranking position for that promoter. This process was repeated for all of the promoters. The thresholds *t* that we tested were 10 and 17822. When using 17822 as the threshold, we only used the cells that had at least 16000 influencers output from the CI algorithm.

## Funding

This research was supported by the National Institutes of Health grant [1R35GM137974 to Z.W.].

## Conflicts of interest

There is no conflict of interest declared.

## Notes

http://dna.cs.miami.edu/scHiMe/

### Competing Interest Statement

The authors have declared no competing interest.

### Summary of Updates

Minor change on Figure 3

