## Supplementary document for "scHiMe: Predicting single-cell DNA methylation levels based on single-cell Hi-C data"

### Supplementary figures

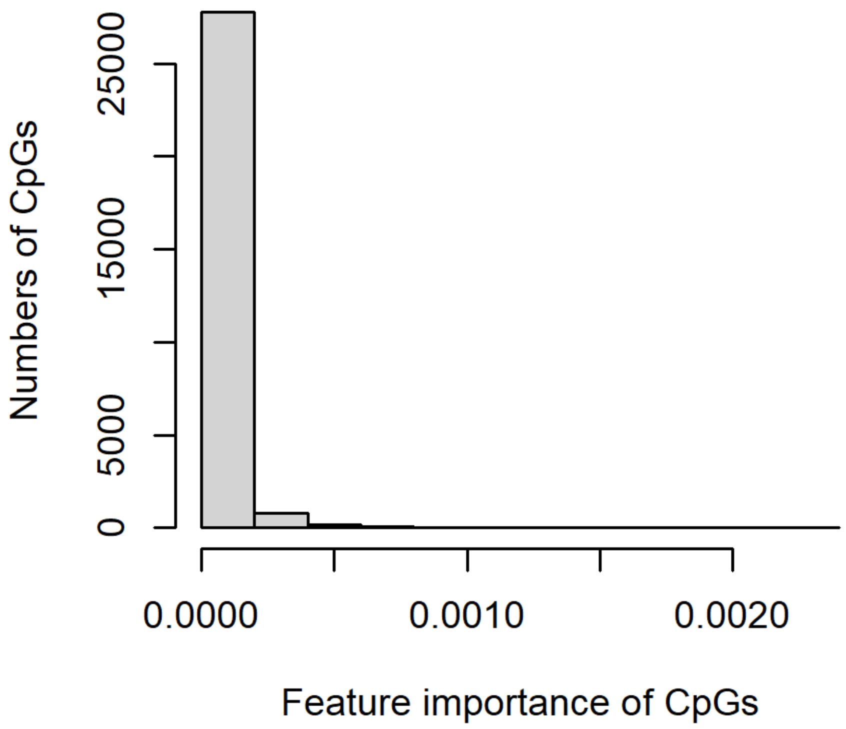


**Fig 1.** Distribution of the number of cytosines and guanines having various values of feature importance.

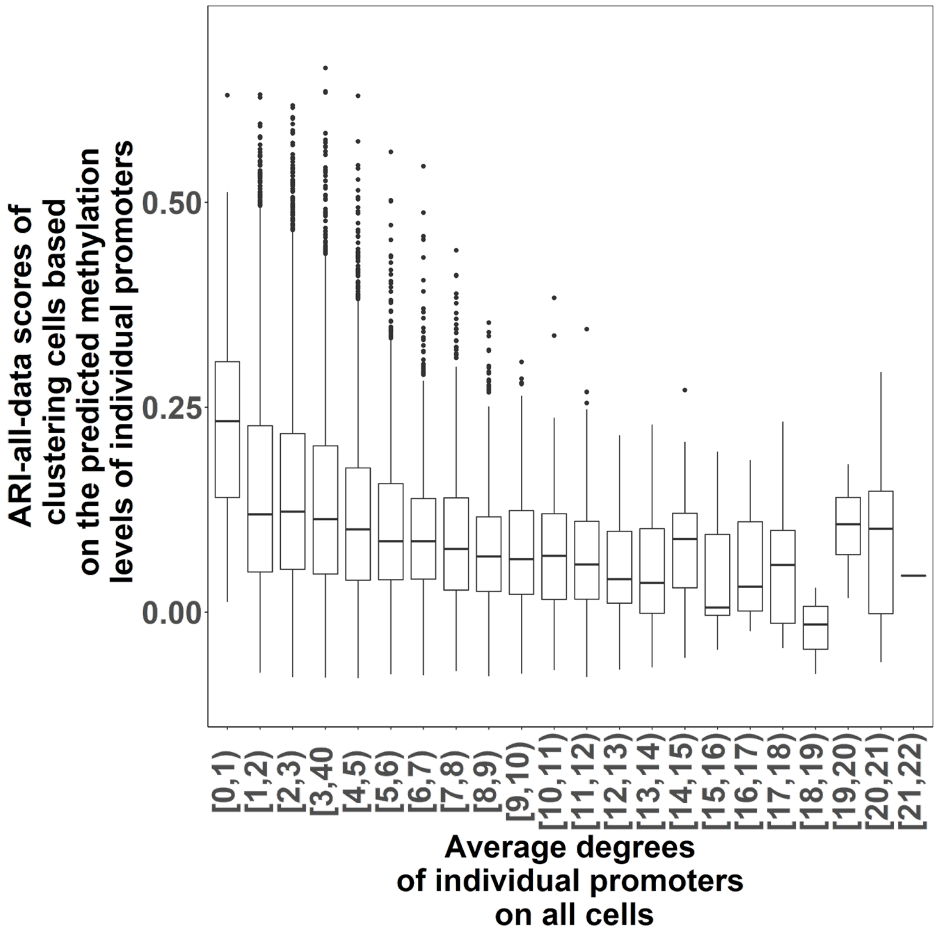


**Fig 2.** ARI-all-data scores of clustering cells based on the predicted methylation levels of the individual promoters of different average degrees.

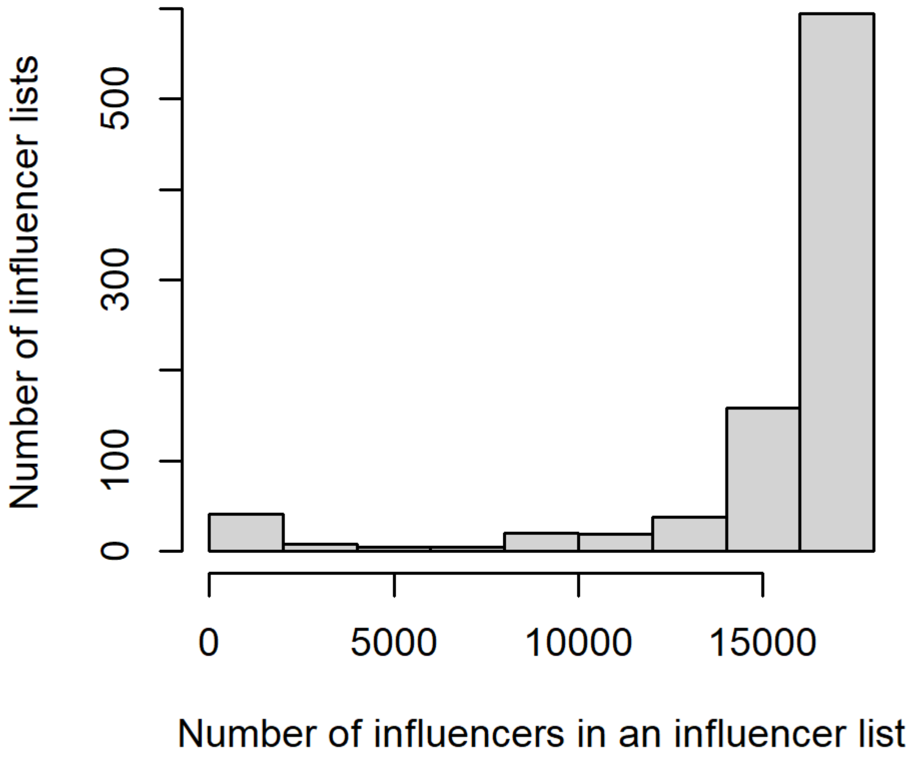


**Fig 3.** The distribution of the number of influencers found for all the cells in data set 2.

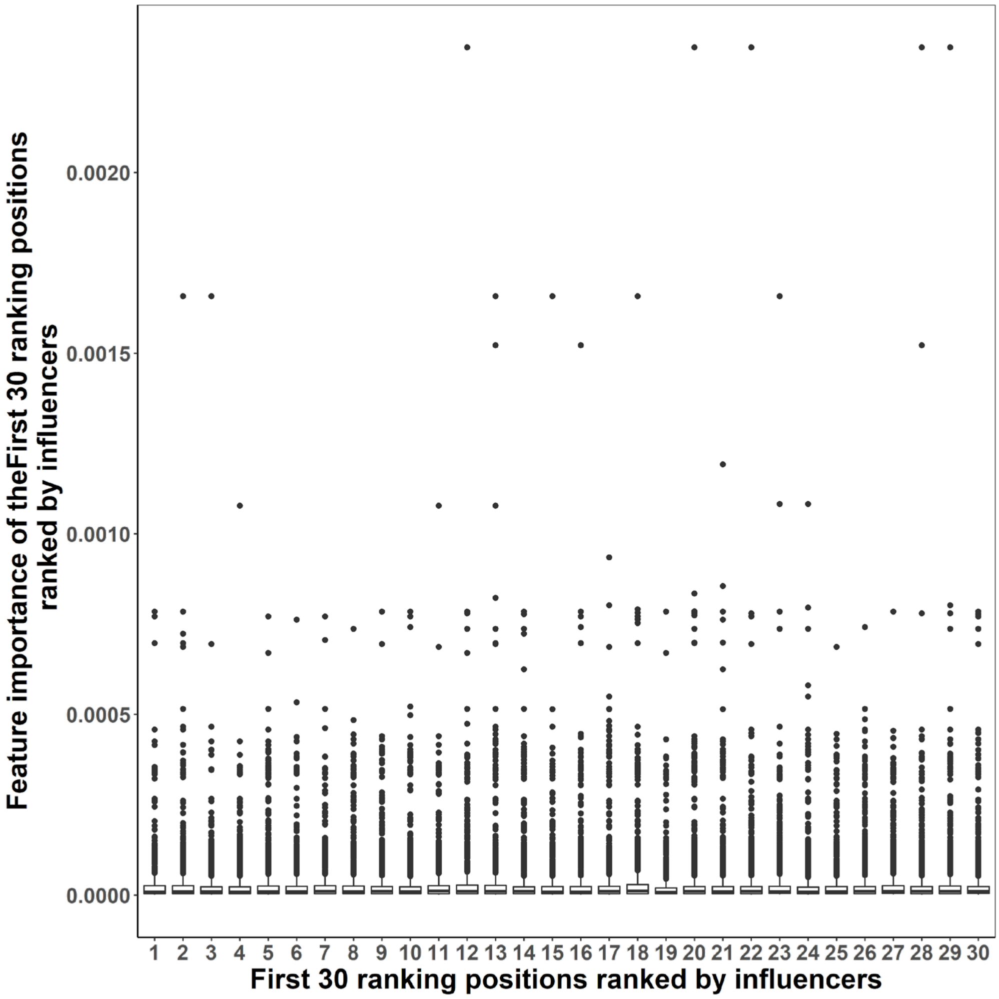


**Fig 4.** The feature importance of the Cs and Gs of CpGs that exist in the promoters at the same ranking positions. The ranking positions are from all the lists of influencers.

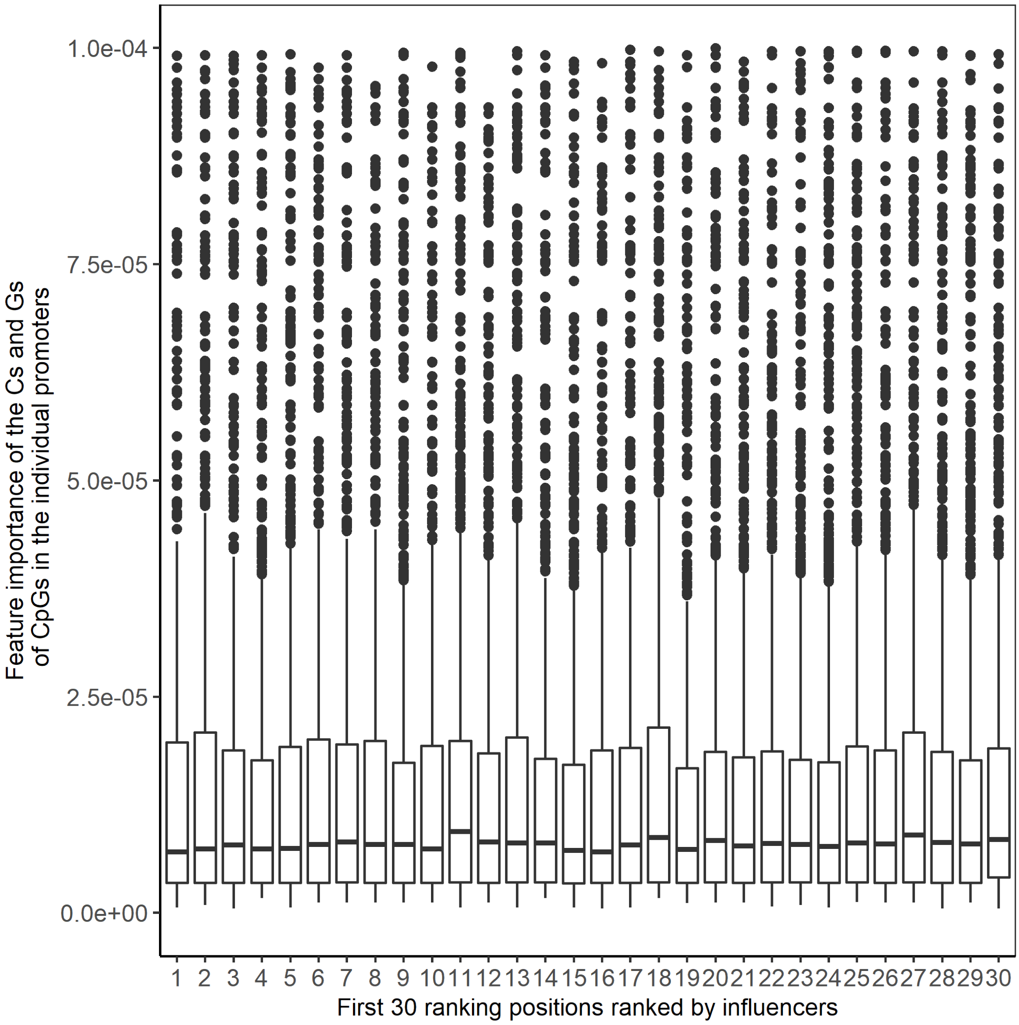


**Fig 5.** The feature importance of the CpGs that exist in the promoters at the same ranking positions from all the lists of influencers. This figure only shows a part of the feature importance from 0 to 0.0001

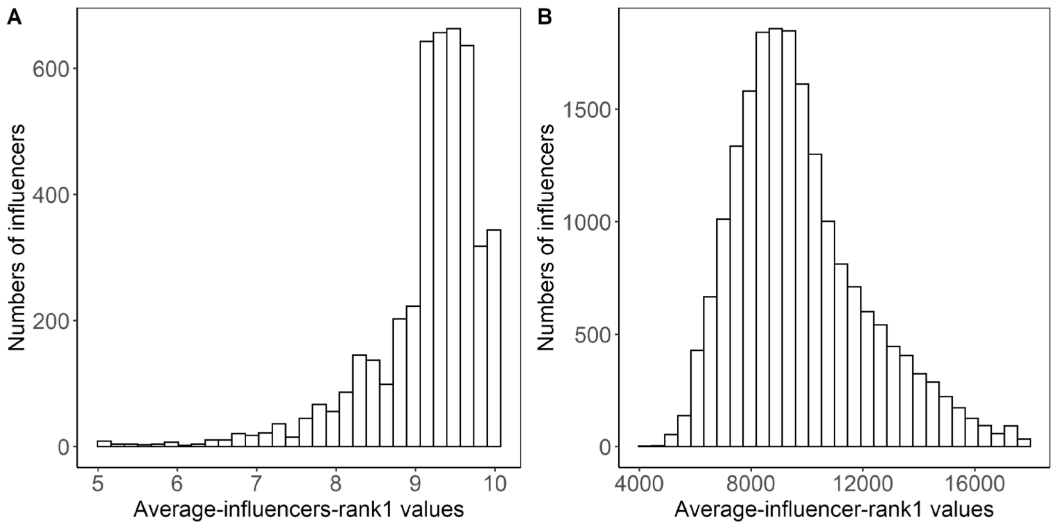


**Fig 6. A-B.** Distribution of the number of influencers with various average-influencer-rank1. **A**. The influencers are top 10 most significant influencers from all cells. **B.** The influencers are from the cells that have at least 16000 influencers detected.

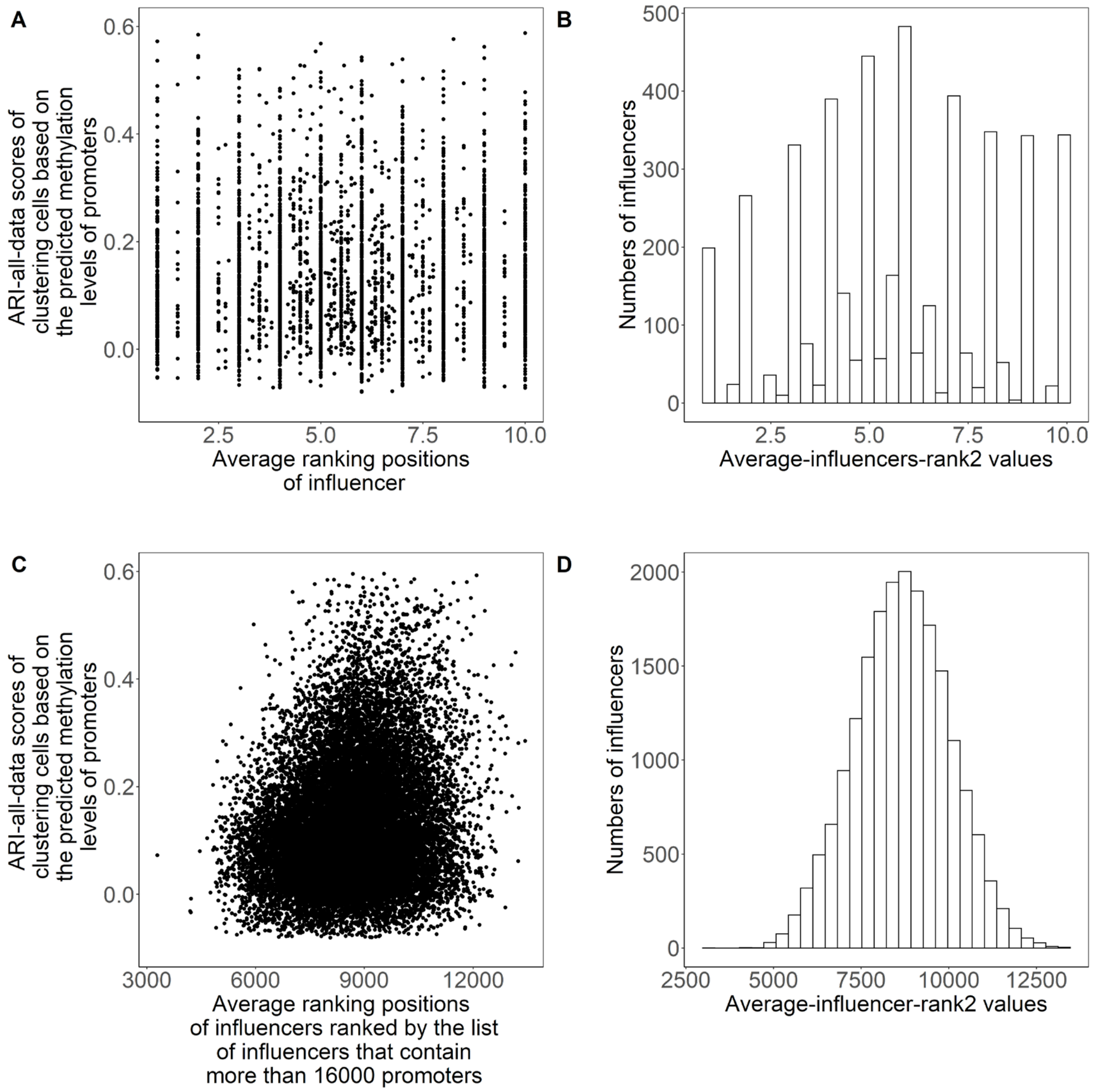


**Fig 7. A.** ARI-all-data scores of clustering cells based on the predicted methylation levels on all cells based on the top 10 most significant influencers (promoters). **B.** Distribution of the number of influencers with average-influencer-rank2. The influencers are top 10 most significant influencers form all cells. **C.** Similar to **A**, the influencers are from the cells that have at least 16000 influencers (promoters) in the list. **D.** Distribution of the number of influencers with average-influencer-rank2. The influencers are top 10 most significant influencers form all cells. The influencers are from the cells that have at least 16000 influencers detected.

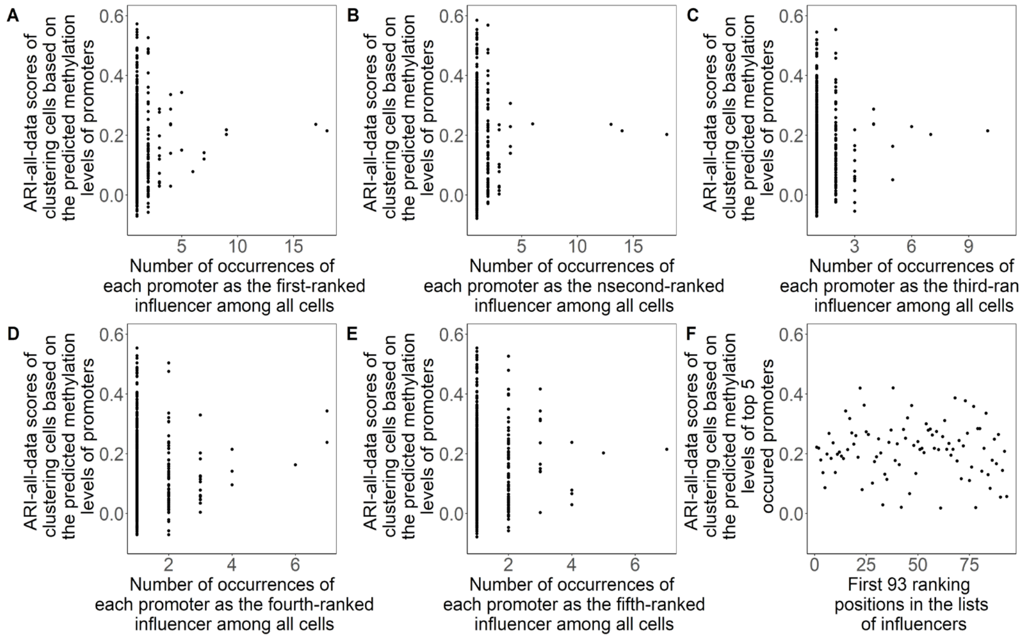


**Fig 8. A-E.** The ARI-all-data scores of clustering cells based on the predicted methylation levels of influencers (promoters) from No. 1 to No. 5 positions in the lists of influencers. **F.** The ARI-all-data scores of clustering cells based on predicted methylation levels of top 5 occurred promoters.

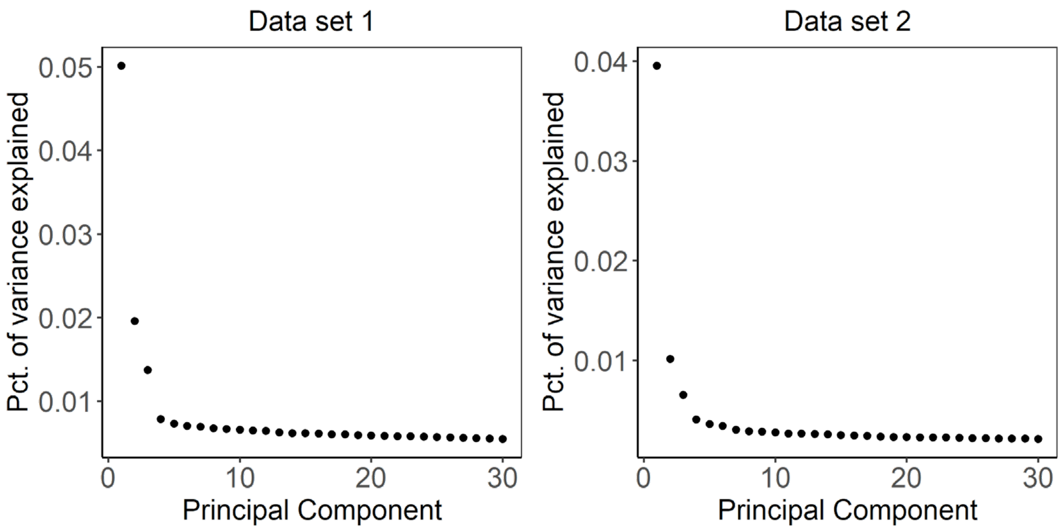


**Fig 9.** The percentage of variance in the data explained by each principal component in the principal component analysis, sorted from highest to lowest.
